# Tendon fibroblast inflammatory responses depend on NF-κβ and JAK/STAT signaling and alter mechanotransduction pathways

**DOI:** 10.1101/2025.07.04.663209

**Authors:** McKenzie Sup, MinKyu M. Kim, Lee Song, Stavros Thomopoulos

**Affiliations:** Department of Biomedical Engineering, Columbia University, New York, NY 10027; Department of Orthopaedic Surgery, Columbia University, New York, NY 10032

**Keywords:** tendinopathy, inflammation, macrophages, JAK/STAT, NF-κβ, mechanotransduction

## Abstract

Tendon pathologies, including both chronic injuries and acute tendon tears, are some of the most common musculoskeletal injuries. Recent studies have suggested the importance of inflammation in the healing process in both acute and chronic tendon injury. However, there remain gaps in knowledge that hinder progress in the development of therapeutics to improve healing. A more complete characterization of the inflammatory response in tendon is needed, by defining the relative roles of different molecular pathways, and determining how these pathways interact with tendon mechanobiology. To investigate these questions, an in vitro model was developed, wherein the complexity of the in vivo healing environment was simulated by applying M1 macrophage conditioned media (M1-CM) to tendon fibroblasts (TFs). Characterization of the M1-CM and its effect on TFs revealed a robust inflammatory response, including upregulation of over 500 genes and increased secretion of several cytokines in TFs. The NF-κβ and JAK/STAT signaling pathways were necessary for the response to M1-CM, and each pathway was responsible for different downstream responses to inflammation in TFs. When considering the role of mechanical loading in tendon responses to inflammation, it was found that TF responses to loading were altered by the presence of an inflammatory stimulus. Analysis of the genes that responded differently to loading with inflammation present suggested changes in pathways involving extracellular matrix organization and G protein signaling. Mathematical modeling based upon these results revealed time-dependent suppression of mechanosensitivity, suggesting that therapeutic timing of inflammatory or anti-inflammatory interventions could restore or attenuate mechanical responsiveness to modulate rehabilitation outcomes. Results reveal that inflammation disrupts mechanosensitivity in tendon healing, and suggest potential pathways for therapeutic intervention.

## Introduction

Tendon and ligament injuries are common medical conditions that result in a high socioeconomic burden. These injuries can occur in several areas of the body, including the shoulder, ankle, knee, and hand, and have varied epidemiology according to the tendon affected ^1^. Achilles tendon injuries, for example, are more prevalent in males than females, and typically occur during sports or moderate to intense physical activity ^2,3^. Supraspinatus tendon tears, meanwhile, are most common in the elderly population, with a prevalence reaching 50% after age 66 ^4^. After a torn tendon heals, its mechanical properties are inferior to the native tissue, leading to reduced functionality ^5^. This is in large part due to the replacement of aligned collagen I with a disorganized scar tissue matrix that forms during the healing process ^6^. As a result, patients often have increased pain, reduced strength, and decreased range of motion even after the tendon healing process is fully complete, which affects their daily activities and athletic performance. Additionally, the impaired mechanical properties greatly increase the risk of retear, a common occurrence in previously ruptured tendons; the rotator cuff, in particular, has re-rupture rates of 20-94% ^7–9^. Surgical techniques for repairing tendons are far from optimal, and in some cases may not significantly improve outcomes compared with non-operative treatment ^10^.

Understanding the tendon healing process is essential for improving interventions. The healing process consists of three main phases: inflammation, proliferation, and remodeling. The inflammatory phase during the first 3-5 days post-injury is characterized by a massive infiltration of immune cells, including neutrophils and macrophages ^11^. In the proliferative stage, lasting up to a few weeks, deposition of scar tissue begins to dominate, and acute inflammation subsides ^12^. The remodeling phase follows, and can persist for many months after the initial injury, resulting in a healed tissue that is ultimately mechanically inferior to the pre-rupture healthy tendon ^5,12^.

The role of inflammatory mediators in tendon healing remains paradoxical, with evidence supporting both beneficial and detrimental effects. This ambiguity is exemplified by interleukin-6 (IL-6), a key cytokine in tendon injury responses. Some studies demonstrate that IL-6 can stimulate collagen synthesis and upregulate extracellular matrix gene expression, potentially supporting tendon repair in the early healing phase ^13–15^. For instance, IL-6 infusion in human peritendinous tissue increases type I collagen synthesis ^14^, and IL-6 upregulates TGF-β1 and ECM genes in tendon progenitor cells ^13^.

Conversely, chronic or excessive IL-6 signaling suppresses type I collagen gene expression and lysyl oxidase, and is associated with features of tendinopathy rather than regeneration ^16,17^. Moreover, while IL-6 promotes proliferation of tendon-derived stem cells, it simultaneously inhibits their tenogenic differentiation, suggesting potential for maladaptive repair if not tightly regulated ^18^. This context-dependent duality extends beyond IL-6 to the broader inflammatory response, where timing, magnitude, and cellular context determine whether inflammation promotes healing or drives pathology ^19–22^.

Given this ambiguity, attempts to improve outcomes by globally suppressing inflammation in the early stages have largely been unsuccessful. However, more targeted approaches that modulate specific inflammatory pathways have shown better results. For example, inhibition of the nuclear factor kappa β (NF-κβ) pathway resulted in improved healing in a rat rotator cuff injury model ^23^. Another targeted approach that has been investigated recently is inhibition of the Janus kinase/signal transducer and activator of transcription (JAK/STAT) pathway; this approach led to increased mechanical properties and collagen content in an *ex vivo* model of tendon injury. ^24^. Despite these encouraging studies, our understanding of major signaling pathways that govern tendon inflammation is still lacking, which hampers the development of disease modifying treatments for tendon injuries. To step towards resolving the fundamental paradox of whether inflammation is beneficial or detrimental, we sought to develop quantitative tools to understand and control the inflammatory response in tendon, including factors such as IL-6 production.

A second promising approach to improve outcomes is to modulate the mechanical loading environment. However, the role of mechanical loading during tendon healing, particularly in the context of the inflammatory environment, is poorly understood. Mechanical loading is necessary for tendon healing, but the cellular-level mechanisms governing this response remain unclear ^25^. Under healthy conditions, exercise regulates production of type I collagen as well as expression and secretion of IL6 ^26–28^.

Because tendon matrix regulation and cytokine production are also affected by the presence of inflammation, there may be interactions between inflammatory and mechanical responses that are important to consider in tendon healing ^12^. Responses to acute inflammation are typically rapid and high in magnitude, and thus may override or suppress typical responses to physiological loading. While few studies have investigated the mechanisms driving crosstalk between inflammatory and mechanical stimuli, evidence suggests that they are integrally connected. For example, patient responses to exercise include inflammation markers such as increased IL-6 production and matrix metalloproteinase (MMP) production ^29,30^. Additionally, pathways that have been shown to be mechanoresponsive *in vitro* have also been implicated in inflammation, including NF-κβ, mitogen-activated protein kinase (MAPK), and JAK/STAT. Finally, previous studies have looked at the effects of inflammation on tendon fibroblast responses to loading *in vitro* ^31,32^. In these studies, the addition of the cytokine IL1β to tendon fibroblasts subjected to loading led to changes in gene expression. Interestingly, low levels of cyclic strain suppressed the IL1β inflammatory response, while high levels of cyclic strain increased it ^31^. An improved understanding of the tendon mechanoresponses under inflammatory conditions will help to guide further research into the use of mechanical loading to treat tendon pathologies, which currently have variable outcomes in the clinic ^7,33–35^.

Characterizing the inflammatory response in tendon, including the role of different signaling pathways and the role of mechanical loading, requires cellular level analysis. Microscale mechanical loading cannot be precisely controlled *in vivo*, and a dose-ranging study of various inhibitors of specific inflammatory pathways would require large numbers of animals. Thus, this study uses an *in vitro* approach wherein the complexity of the *in vivo* healing environment is simulated by the presence of signaling from M1 macrophages, since this type of macrophage is highly pro-inflammatory and is the predominant cell type in early stages of tendon healing (in contrast to the anti-inflammatory M2 macrophage which is predominant in later stages of healing) ^36^. We hypothesized that M1 macrophage paracrine signaling would induce a widespread pro-inflammatory response in tendon cells, and that both the NF-κβ pathway and JAK/STAT pathways would be necessary in this response. Additionally, we hypothesized that mechanical loading responses would be altered in the inflammatory environment; typical responses would be suppressed and loading would mitigate the inflammatory phenotype.

## Methods

### Animal information

Primary tendon fibroblast (TF) and macrophage cultures were established from cells isolated from 2-month-old C57BL/6 mice. Housing, breeding, and the use of mice were performed in accordance with the Institutional Animal Care and Use Committee at Columbia University. Tendon fibroblast were isolated from tail tendons and macrophages were derived from bone marrow (Figure 1).

**Figure 1:**
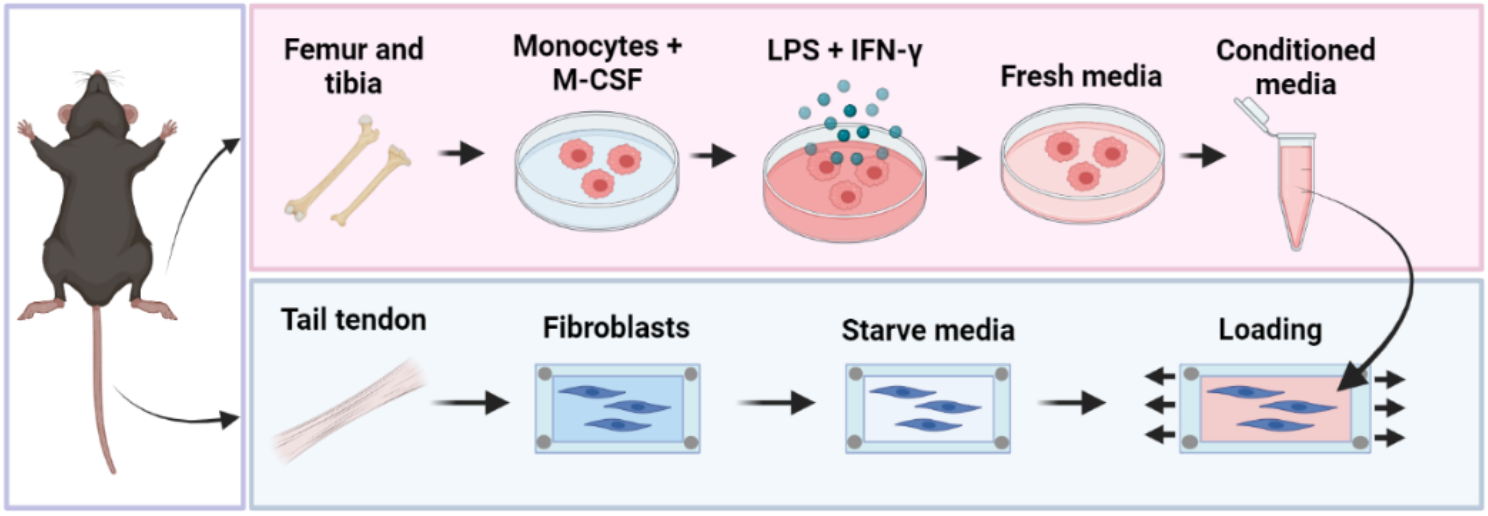
Methods overview.

### Tendon fibroblast isolation and culture

TFs were isolated from tail tendon by digestion in 0.2% collagenase type II (Worthington Biochemical) in 5mL alpha-modified Eagle’s medium (Alpha-mem) in a shaking incubator at 270 rpm and 37 degrees C for 1 hour, or until no intact tissue pieces remained. Tissue was vortexed every 30 min during digestion. Then, cells were strained through a 100uM cell strainer over a sterile 50 mL falcon tube, pelleted by centrifugation at 450 g for 5 min, and resuspended in Alpha-MEM with 10% fetal bovine serum (FBS) and 1% penicillin/streptomycin (P/S). Media was changed every 2 days and cells were passaged at 80% confluence. All TFs were seeded for experiments at passage 2 or 3, and were serum starved at 1% FBS for 24 hours prior to stimulation with M1-CM, application of loading, and/or addition of inhibitor.

### Macrophage isolation and culture

Bone marrow aspirate was collected from femur and tibia, then added to 60-mm Petri dishes in 80% Roswell Park Memorial Institute (RPMI) medium (with 10% FBS and 1% P/S), and 20% L929 cell-conditioned medium as a source of macrophage colony stimulating factor (M-CSF). Fresh media was added to cultures every other day over 5 days to culture macrophages to confluence. Then, macrophages were polarized to a pro-inflammatory M1 phenotype by stimulation with 100ng/mL lipopolysaccharides (LPS) (Sigma Aldrich, #L4391-1MG) and 20ng/mL interferon gamma (IFN-γ) (PeproTech, #315-05) for 24 hours.

### M1 conditioned media collection and application to TFs

After 24-hour polarization, M1-stimulation media was removed from the macrophages, two phosphate-buffer-saline (PBS) rinses were performed, and fresh macrophage culture media was applied. 24 hours later, the supernatant was collected from the activated macrophages, sterile filtered with a 0.2 uM filter, and then added to the TFs in a 1:1 ratio with fresh TF culture media (N=4, 2 male and 2 female mice).

### Mechanical loading

Polydimethylsiloxane (PDMS) stretch chambers (STB-CH-04, STREX Co., Osaka, Japan) with a surface area of 4cm^2^ were autoclaved, coated with 50 mg/mL fibronectin (EMD Millipore, #FC010) for 1hr, and finally rinsed twice with PBS for 5 minutes. TFs from N=4 (2 male, 2 female) mice, grown to confluence at passage 1, were then lifted using Trypsin-EDTA 0.05% (Thermo Scientific, #25300120), and seeded into each chamber in 2mL of Alpha-MEM at a density of 2500 cells/ cm^2^. Cells were returned to the incubator and allowed to attach for 24 hours prior to the start of loading experiments. On the day of loading, half of the chambers received M1-CM, and the other half macrophage culture media. Half of the M1-CM-treated chambers and half of the untreated chambers were then placed into the STB-1400 STREX cell stretch system such that they were held at gauge length prior to onset of loading. Four total chambers (one for each treatment group) were used per biological replicate. The remaining chambers were similarly manipulated and then placed in petri dishes inside the same incubator. All chambers were then left to rest for 1 hr after loading was set up before strain was applied. Cyclical loading was initiated at a frequency of 0.5 Hz and strain level of 8% and maintained for 24 hrs. Strain and frequency levels were set on the STB-1400 according to manufacturer specifications, and were selected based on levels previously established to be in the physiological range for tendon ^37^.

### NF-κβ and JAK/STAT inhibition

For inhibition experiments, TFs were plated in a 12-well plate at a density of 1.7 × 10^5^ cells per well with 1mL of culture medium. IKK-2 Inhibitor VIII, an inhibitor of IKK-2 activity, was used to suppress activation of the NF-κβ pathway (Sigma-Aldrich, #401487). IKK-2 is crucial in formation of the IKK complex that induces degradation of the IkB subunit, allowing NF-κβ to translocate into the nucleus. IKK-2 was added to TFs at a range of concentrations from 1-10uM, with or without the simultaneous application of M1-CM, for a period of 24 hours before analysis of protein and gene expression. Ruxolitinib (MedChem Express, #HY-50856) was used to inhibit JAK/STAT signaling. Ruxolitinib is an inhibitor of JAK1 and JAK2, which are crucial for the phosphorylation of STAT and activation of the JAK/STAT signaling pathway. Ruxolitinib was added to TFs at a range of concentrations from 1-10uM, with or without the simultaneous application of M1-CM, for a period of 24 hours before analysis of protein and gene expression. There were 4 biological replicates (N=4) for inhibitor studies, with 2 male and 2 female mice. TFs and macrophages were isolated from each biological replicate.

### Gene expression analysis

For loading experiments, immediately after the end of the loading period, PDMS chambers were removed from the STB-1400 STREX cell stretch system and placed into petri dishes. Chambers were rinsed twice with sterile room temperature PBS, treated with 350μL of RNeasy Mini Kit (Qiagen) RLT buffer, and scraped with a P-1000 pipet tip to facilitate removal of cells from PDMS surface. Finally, cell lysate was removed from chambers, mixed with equal volume molecular biology grade 70% ethanol, and added to RNeasy Mini Kit spin columns (Qiagen), before RNA purification was conducted according to manufacturer protocol. For inhibitor experiments, RNA isolation was performed similarly, but on 12-well tissue culture plates. For RNA designated for qPCR, quality and quantity was checked using a plate reader, and 100 ng of RNA for each sample was reverse transcribed to cDNA using the High-Capacity cDNA Reverse Transcription Kit (Applied Biosystems). qRT-PCR was conducted in the QuantStudio 6 Flex (Applied Biosystems) using PowerUp SYBR Green Master Mix (Applied Biosystems). For TF Response to M1 CM Treatment, gene expression was normalized to GAPDH and then expressed as fold change of M1-CM-treated gene expression relative to untreated control cells. For loading response, gene expression was normalized to GAPDH and expressed as fold change of loaded cells relative to unloaded cells, separately for the M1-CM-treated and untreated groups. For inhibition responses, gene expression of each inhibition group was expressed as a quantity relative to GAPDH expression in each sample. The complete set of genes assessed were *Acta2, Ccl2, Col1a, Cxcl1, Cxcl10, Ift88, Il6, Mmp13, Piezo1, Ptgs2, Scx, Sod3*, and *Tgfb*. All primer sequences are listed in Table S1.

### RNA sequencing

RNA intended for RNA sequencing was quantified and checked for quality using the Bioanalyzer (Agilent) through the Columbia University Molecular Pathology core service. Bulk RNA sequencing was performed using the Illumina NovaSeq Platform with TruSeq library prep (depth of > 16M reads per sample) at the Columbia Genome Center. A DESeq2 Output was performed using the Bioconductor DESeq2 package in R with three comparisons: Unloaded Control TFs vs. Loaded Control TFs, Unloaded M1-CM-Treated TFs vs. Loaded M1-CM-Treated TFs, and Unloaded Control TFs vs Unloaded M1-CM-treated TFs. For analysis of M1-CM-treated vs untreated TFs, the threshold for significance was set to p_adj_ <0.05 and log_2_FoldChange > 1.5 or < -1.5, for upregulation or downregulation, respectively. For analysis of loaded vs unloaded TFs, the threshold for significance was the same, except that the threshold magnitude for log_2_FoldChange was lowered to 1, because the fold change responses to mechanical loading were observed to be overall lower in magnitude than inflammatory responses. Volcano plots were generated using the tidyverse package and ggplot in R, and Heatmaps were generated using the ComplexHeatmap package (Gu, Z. Complex Heatmap Visualization. IMETA 2022). For analysis of TF response to M1-CM, KEGG enrichment analysis was performed on an order ranked gene list with an adjusted p value of <0.05 using the gseKEGG function in the ClusterProfiler package in R. For analysis of loading response, Reactome Pathway Analysis was done using the ReactomePA package in R, which used a non-order ranked gene list as input. This method was chosen in favor of the ranked list used for analysis of inflammatory responses alone, since mechanoresponses were generally smaller in magnitude than inflammatory ones, and the goal was to identify possible interactions between the responses rather than magnitude of their interactions. The gene counts for each experimental group were analyzed for significant differences in expression of transcripts using the Bioconductor DESeq2 package in R. Heatmaps were generated using the ComplexHeatmap package (Gu, Z. Complex Heatmap Visualization. iMeta 2022). Volcano plots were generated using the tidyverse package in R. The ggplot2 package and AnnotationDBI packages in R were used for both analyses.

### *LegendPLEX* multiplex immunoassay

Cell supernatant was collected 24 hours after M1-CM stimulation, and then immediately frozen at –80°C until assay was performed. On the day of the assay, supernatant samples were thawed, centrifugation was performed for 5 minutes at 12000g to collect any remaining cell debris, and finally the samples were transferred to the 96-well plate provided in the mouse anti-virus response panel LegendPLEX kit. This pre-defined panel included 13 proteins: IFNγ, TNFα, IL1β, CXCL1, CCL2, CCL5, CXCL10, IL10, IFNβ, IFNα, IL12p70, GMCSF, and IL6. All steps were performed according to manufacturer instructions. There were 3 biological replicates (N=1 male and N=2 females), and TFs and macrophages were isolated from each biological replicate.

### Cytotoxicity Assays

For LIVE/DEAD imaging, after 24h treatment with M1-CM and or inhibitor, cell supernatant was removed and cells were washed with warm PBS. Working solution of Live Green and Dead Red components were added to cells from the LIVE/DEAD Cell Imaging Kit (ThermoFisher) according to manufacturer instructions. For qualitative confirmation of the observed differences in cell confluence, one biological replicate (N=1 male) and N=2 technical replicates were included. Images at five different field of views were taken at 10x magnification per experimental group, and one representative image from each group was selected and shown. For the LDH assay (MilliporeSigma), cell supernatant was removed from TFs from N=1 female and N=2 males after 24h treatment with M1-CM and/or inhibitor, and frozen at –80°C until assay was performed. On the day of the assay, supernatant samples were thawed, centrifugation was performed for 5 minutes at 12000g to collect any remaining cell debris, and finally the samples were transferred a 96-well plate. LDH concentration was calculated using a standard curve, and differences in LDH concentration in supernatants were compared between M1-CM treatment only, and groups with inhibition + M1-CM. Experimental groups with more than one sample not within the detection limit of the standard curve (calculated as negative concentration) were not included in statistical analysis.

### Western Blot

Cells were plated in 6-well plates at a density of 7 × 10^5^ in Alpha-mem with 10% FBS, and then serum starved at 1% FBS Alpha-mem for 24 before inhibitors and/or M1-CM were added. After 24 hours treatment with inhibitor or positive control, cells were lysed with radioimmunoprecipitation assay lysis buffer (Thermo Fisher Scientific) supplemented with protease and phosphatase inhibitors (Thermo Fisher Scientific #78441). Total protein was quantified with the Pierce BCA Protein Assay Kit (Thermo Fisher Scientific #23227). Equal amounts of total cell lysate (60ug) were subjected to SDS-electrophoresis on 10% bis-acrylamide gels. Gels were run at a constant voltage of 100mV for 1 hour (PowerPac 3000, Bio-Rad). Membrane transfer was performed at constant 100V for 1 hour. Blots were treated with blocking solution (5% nonfat milk in TBS Tween 20) for 1 hour prior to application of each primary antibody. Immunoblotting was completed for Phospho-p65, p65, and GAPDH for IKK-2 inhibitor validation, each at a concentration of 1:1000 in blocking solution (Cell Signaling #3033T, #8242T, and #5174T, respectively). Immunoblotting for Ruxolitinib validation was completed for Phospho-STAT1, STAT1, and GAPDH, each at a concentration of 1:1000 in blocking solution (Cell Signaling #9167T, #14994T, and #5174T, respectively). Secondary antibody, Anti-Rabbit IgG HRP-linked (Cell Signaling #7074s), was used at 1:2000 in 5% nonfat milk in TBS Tween 20. A Li-Cor Odyssey scanner was used to visualize immunoblots.

### Statistics

For RNA sequencing analysis, statistical significance was determined using the DESeq2, ClusterProfiler, and AnnotationDBI packages referenced above. For qPCR validation of M1-CM-treatment RNAsequencing results, paired t tests were performed for each gene between control TFs and TFs treated with M1 CM. For analysis of proteins secreted by M1 macrophages, paired t tests were performed for each protein between M1 CM and M0 CM, to assess which proteins were significantly up-regulated by pro-inflammatory polarization. To determine the TF-specific cytokine secretion profile compared to M1 cytokine secretion, a one-way ANOVA was performed to test for the effect of cell treatment group on cytokine concentrations, with treatment groups being control TFs, M1 CM, and TFs treated with M1 CM. Holm-Sidak adjustment for multiple comparisons was performed to determine significance of differences between each group. For loading experiments, paired t-tests were performed to assess for differences between loaded TFs and unloaded TFs, with unloaded controls present in each group (M1-CM treated and untreated). Delta CT values normalized to GAPDH were used for statistical testing in all gene expression analysis. For inhibition experiments, a one-way ANOVA was performed for each gene to assess for an effect of inhibition treatment on gene expression in TFs treated with M1-CM, followed by Holm-Sidak adjustment for multiple comparisons between TFs treated with each inhibitor.

### Mathematical Chemomechanical Model

We developed a simplified mathematical model to capture the key finding that mechanical loading responses are suppressed under inflammatory conditions. The model uses ordinary differential equations (ODEs) to describe the temporal dynamics of two key variables: inflammatory factors (*C*_*str*_) and stress response factors (*C*_*inf*_). The inflammatory factor variable represents the combined effects of NF-κB and JAK/STAT pathway activation in response to M1-CM stimulation. This lumped parameter captures the net inflammatory state of the system. The stress response variable represents mechanically-induced factors such as IL-6 and MMP-13, which are normally upregulated by loading but suppressed under inflammatory conditions.

The kinetics of the inflammatory cytokines follow:

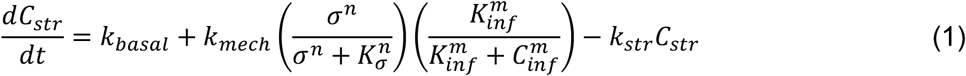

and

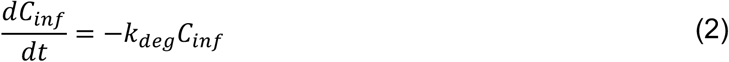

Where *σ* is a normalized mechanical loading magnitude; *k*_*basal*_, *k*_*mech*_, *k*_*str*_, and *k*_*deg*_ are rate constants; *K*_*inf*_ and *K*_*σ*_represent loading levels for half-maximal mechanical activation and the inflammatory level for half-maximal suppression, respectively; and the Hill coefficients *n* and *m* capture the cooperativity of these responses.

The first equation describes the decay of inflammatory factors over time, representing the clearance of M1-CM components. The second equation captures three key processes: basal production (*k*_*basal*_), which represents constitutive expression of stress factors; mechanical induction with inflammatory suppression, where the term 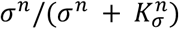 represents mechanical activation following a Hill function, while 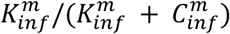 represents inflammatory suppression; and degradation(*k*_*str*_*C*_*str*_),which represents first-order decay of stress factors. When *C*_*inf*_ is low, the suppression term approaches 1 (no suppression), and when *C*_*inf*_ is high, this term approaches 0 (complete suppression).

Parameters were chosen to reproduce experimental observations based upon ranges found in the literature, with *k*_*deg*_ = 10^–1^ h^−1^ (inflammatory factor half-life ∼7 hours), *k*_*basal*_ = 10^–1^ h^−1^, k_mech = 2.0 (20-fold maximum induction by loading), *k*_*str*_ = 0.5 h^−1^ (stress factor half-life ∼1.4 hours), *K*_*σ*_= 0.2 (half-maximal activation at applied loading level), *K*_*inf*_ = 0.5 (half-maximal suppression at moderate inflammation), and *n* = *m* = 2 (cooperative responses) ^38–43^. For M1-CM treated conditions, the initial inflammatory factor concentration was set to *C*_*inf*_(0) = 2.0 (normalized units), while for control conditions *C*_*inf*_0) = 0. For both conditions, the initial stress factor level was set to *C*_*str*_(0) = 0.2 (baseline level). The model was solved numerically using MATLAB’s ode45 function.

Key outputs demonstrated that under control conditions (*C*_*inf*_= 0), mechanical loading (*σ*= 0.2) induced a ∼5-fold increase in str, while under inflammatory conditions (*C*_*inf*_ (0) = 2.0), the same mechanical loading produced minimal changes in str due to inflammatory suppression. The inflammatory factor decayed over time while maintaining suppression of mechanically-induced responses throughout the 24-hour period. This simplified model successfully captures the experimental observation that inflammation suppresses normal mechanical responses in tendon fibroblasts, providing a framework for understanding the interplay between inflammatory and mechanical signaling in tendon healing.

## Results

### M1 macrophage media induced a widespread pro-inflammatory response in tendon fibroblasts

To characterize the response of TFs to M1-CM, bulk RNA sequencing analysis was performed on TFs treated with M1-CM versus non-treated control TFs. 550 genes were significantly upregulated, while only 98 genes were significantly downregulated (Figure 2B). A heatmap of the top 50 differentially expressed genes (DEGs) after stimulation of TFs with M1-CM revealed significant upregulation of many genes related to inflammatory processes, including *Irf1*, a master activator of immune responses, and *Stat2*, a signal transducer of the JAK/STAT pathway [24,25] (Figure 2A). Additionally, the chemokines *Ccl2, Ccl7*, and *Cxcl1* were among the top 50 significantly DEGs. Upregulation of *Sod3* and *Socs3* was also observed, suggesting that the TFs were increasing protective mechanisms against inflammation in response to M1-CM ^44,45^. Gene set enrichment and KEGG pathway analysis were both performed to identify upregulated and downregulated pathways, respectively (Figure 2C). Some responses of note were downregulation of metabolism-related gene sets, upregulation of the innate immune response, and upregulation of specific pathways including IL-17, NF-κβ, JAK/STAT, and TNF. qPCR was also performed on a subset of genes found to be robustly upregulated in the RNA sequencing results, confirming the upregulation of *Il6, Mmp13, Cxcl1, Cxcl10, Ccl2*, and *Sod3* (Figure 3).

**Figure 2:**
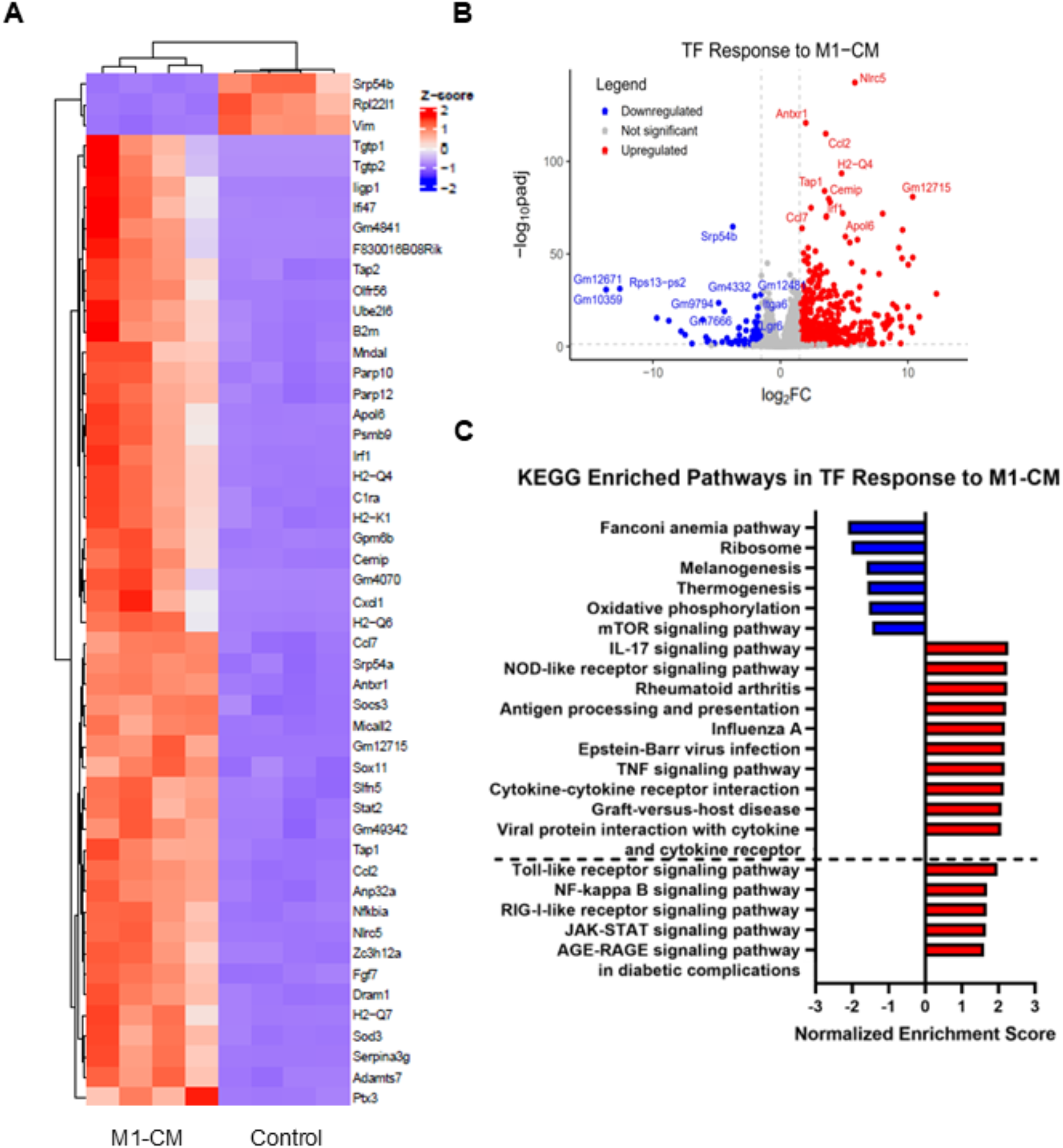
A) Heatmap of top 50 DEGs. B) Volcano plot of significant DEGs. C) KEGG enriched pathways for all downregulated pathways as well as the top 10 and selected additional upregulated pathways.

**Figure 3:**
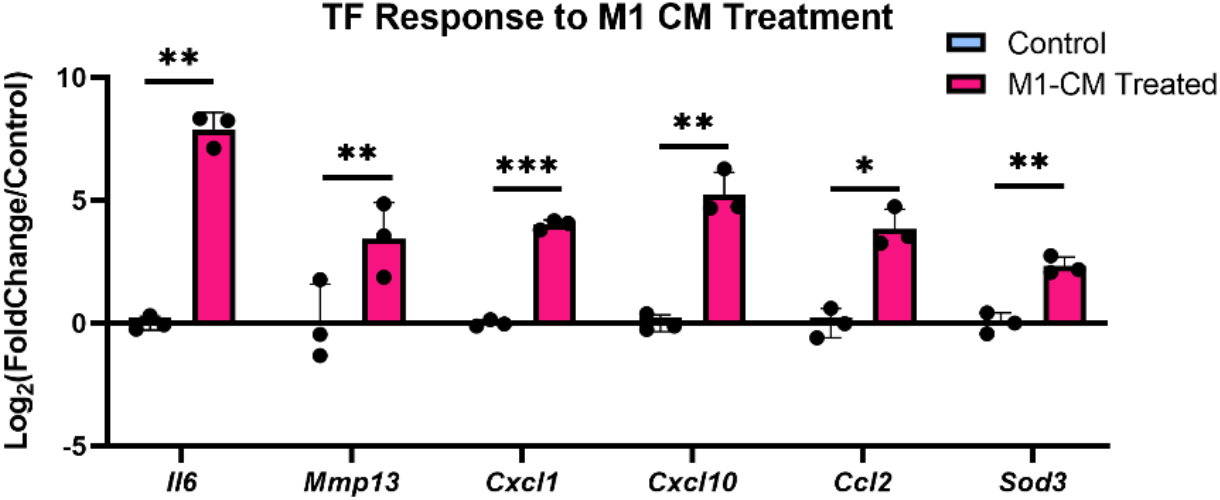
Gene expression based on RT-qPCR of selected genes of interest. Statistical significance was calculated with a paired student’s t test. * p<0.05, ** p<0.01, ** p<0.001, *** p<0.0001. Data shown as mean +/- SD, with individual points representing biologically independent samples.

In response to treatment with M1-CM, TFs contributed a distinct profile of cytokines to the supernatant, in comparison to M1 macrophages alone (Figure 4). For example, TFs did not secrete IFN-y, as the level of this cytokine was lower after 24 hours of incubation with M1-CM than it was initially (i.e., in the M1-CM itself). Levels of IFNβ, IL1β, TNFα, CXCL10, and CCL5 were all higher after M1-CM treatment of TFs compared to the initial levels in M1-CM alone, indicating that TFs were secreting significant amounts of these cytokines. Levels of IL6, Cxcl1, and Ccl2 were not produced at high levels by M1 macrophages (i.e., in M1-CM alone) compared to control TF supernatant, but were produced at very high levels in M1-CM-treated TFs. This indicates that M1 macrophages do not secrete these cytokines in significant quantities, but that TFs stimulated by them do (i.e., these cytokines are tendon-specific).

**Figure 4:**
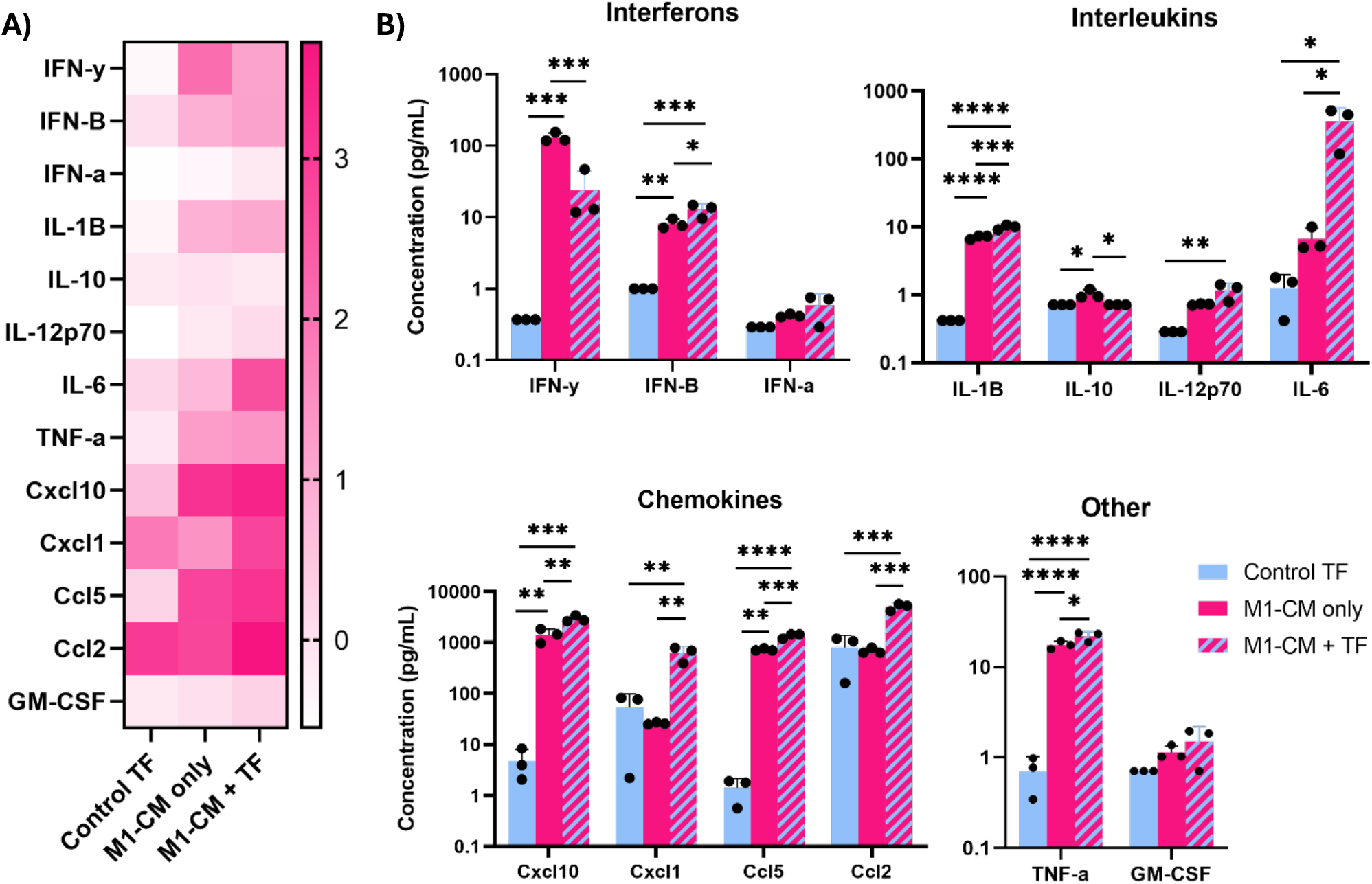
A) Heatmap of log-transformed protein concentrations of each protein from each treatment group. B) Concentrations of each protein concentration from each treatment group, plotted on a log scale. Data is shown as mean +/- SD, with individual points representing biologically independent samples. Statistical significance was calculated using a one-way ANOVA with Holm-Sidak adjustment for multiple comparisons between all treatment groups. * indicates p<0.05, ** p<0.01, *** p<0.001, **** p<0.0001.

### Fibroblasts under M1-CM-induced inflammatory conditions responded differently to loading compared to untreated controls

Mechanical loading is known to regulate many aspects of tendon development and homeostasis, and more recently, has been shown to play a role in healing ^25^. Therefore, we investigated whether the inflammatory activation of TFs in response to M1-CM would affect how they responded to mechanical loading. First, the response to loading of a preliminary panel of genes in both untreated and M1-CM-treated TFs was assessed by qPCR. There was a significant effect of 24 hrs of cyclic tensile loading on TF gene expression (Figure 5). In untreated TFs, *Il6, Mmp13*, and *Ptgs2* were upregulated in response to loading, while *Scx, Acta2*, and *Ift88* were downregulated. Interestingly, M1-CM significantly affected TF responses to loading, dramatically decreasing *Il6* and *Mmp13* responses to loading. Expression of *Scx, Acta2*, and *Ift88* were downregulated due to loading under M1-CM culture conditions, similar to control culture conditions. These results imply an altered mechanoresponses under inflammatory conditions, although the mechanism was unclear and only a subset of genes were affected. Therefore, to comprehensively explore the effect of inflammation on mechanotransduction, RNA sequencing was performed.

**Figure 5:**
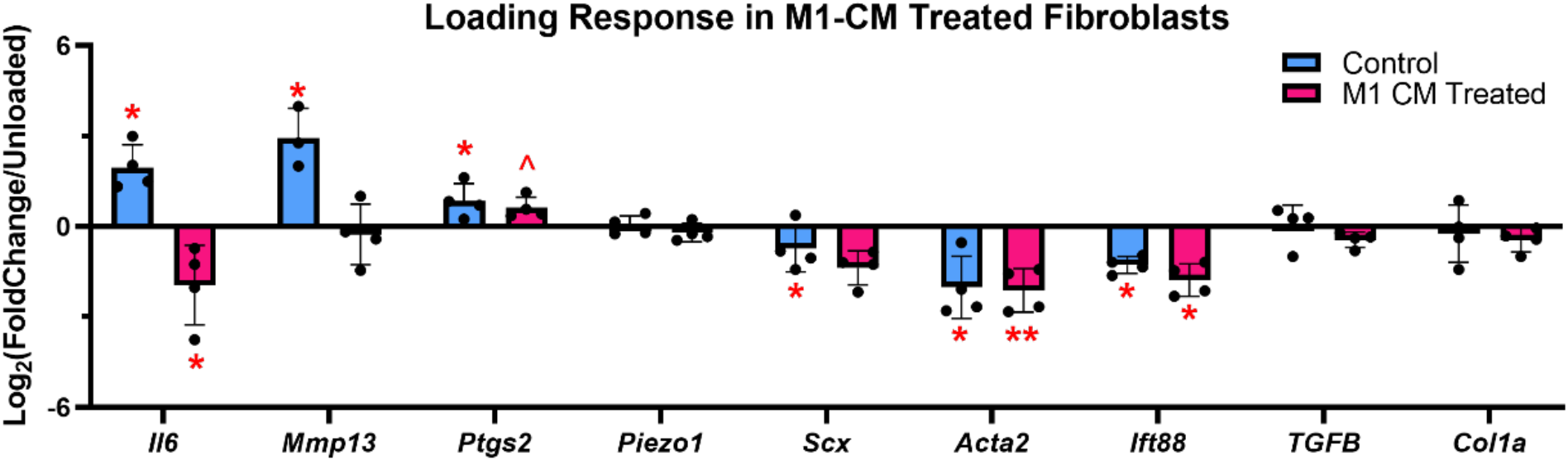
RT-qPCR of select genes related to mechanoresponsiveness, inflammation, and tendon function in response to loading in both treatment groups, compared to unloaded controls in each group. Data is shown as mean +/- SD, with individual points representing biologically independent samples. Statistical significance was calculated using a paired student’s t test. *indicates p<0.05, **indicates p<0.01.

Four groups were analyzed in RNA sequencing: untreated control (UC), M1-CM-treated control (MC), untreated loaded (UL), and M1-CM-treated loaded (ML). The PCA plot generated from the sequencing analysis across all four groups showed distinct clustering of each group (Figure 6C). In TFs treated with M1-CM, 463 genes were upregulated in response to loading, while 1315 genes were downregulated. By comparison, in untreated TFs, 511 genes were upregulated in response to loading, while 1034 genes were downregulated. To determine how much overlap there was in the gene expression responses to loading between the treatment groups, a Venn analysis was generated based on the genes that met the thresholds for significance in each group for upregulation and downregulation by loading (Figure 6E). We identified 650 genes which were downregulated by loading in both treatment groups, while 166 were upregulated in both. Next, because we were primarily interested in the genes that responded differently to loading, we chose to focus our analysis on the genes that did not overlap in the Venn analysis, which consisted of four distinct groups: 1) 345 upregulated by loading only under control conditions; 2) 297 upregulated only under inflammatory conditions; 3) 384 downregulated only under control conditions; and 4) 665 downregulated under inflammatory conditions. However, before proceeding with analysis on these subgroups of interest, we first applied an additional filter, because some of these genes may have just barely missed the criteria for significance in the contrasting treatment group. For example, if a gene went up by 1.1 log_2_FoldChange in control conditions but only 0.9 log_2_FoldChange under inflammatory conditions, simple Venn analysis would group the gene as uniquely regulated by loading depending on the treatment condition, since the gene just barely missed the threshold log_2_FoldChange of 1. Therefore, we adjusted the analysis such that for a gene to be considered differentially regulated by loading depending on treatment condition, the log_2_FoldChange loading response had to have a log_2_FoldChange of the opposite sign in the other treatment group. For example, in order for a gene to be considered upregulated by loading only under control conditions and not under M1-CM-treatment conditions, the gene was required to have a negative log_2_FoldChange response to loading in the M1-CM-treated group. After this additional filtration step, 268 genes remained that met the criteria for being upregulated by loading only under control conditions, while 223 genes were upregulated only under inflammatory conditions (Figure 6E). 70 genes were downregulated only under control conditions, while 396 were downregulated only under inflammatory conditions. These four groups of genes were then pooled together and used for Reactome Pathway Analysis, to determine if specific pathways were responsible for the differential responses to loading (Figure 6F). 12 pathways were identified that met an adjusted statistical significance threshold of p < 0.05, which included extracellular matrix organization, G alpha (i) signaling events, and alternative complement activation Differences in extracellular matrix organization were primarily in matrix metalloproteinase genes (*Mmps)*, and genes for various collagens. For example, *Mmp3, Mmp13*, and *Mmp12* were all upregulated by loading under untreated conditions, but unchanged by loading when M1-CM was present. Meanwhile, *Col15a1. Col3a1*, and *Col12a1* were all unchanged by loading in untreated conditions, but downregulated by loading when M1-CM was present. Genes related to G alpha (i) signalling events included *Npy*, which activates the MAPK pathway and was upregulated by loading under untreated conditions, and *Ednra*, which activates calcium signalling and was downregulated by loading with M1-CM present.

**Figure 6:**
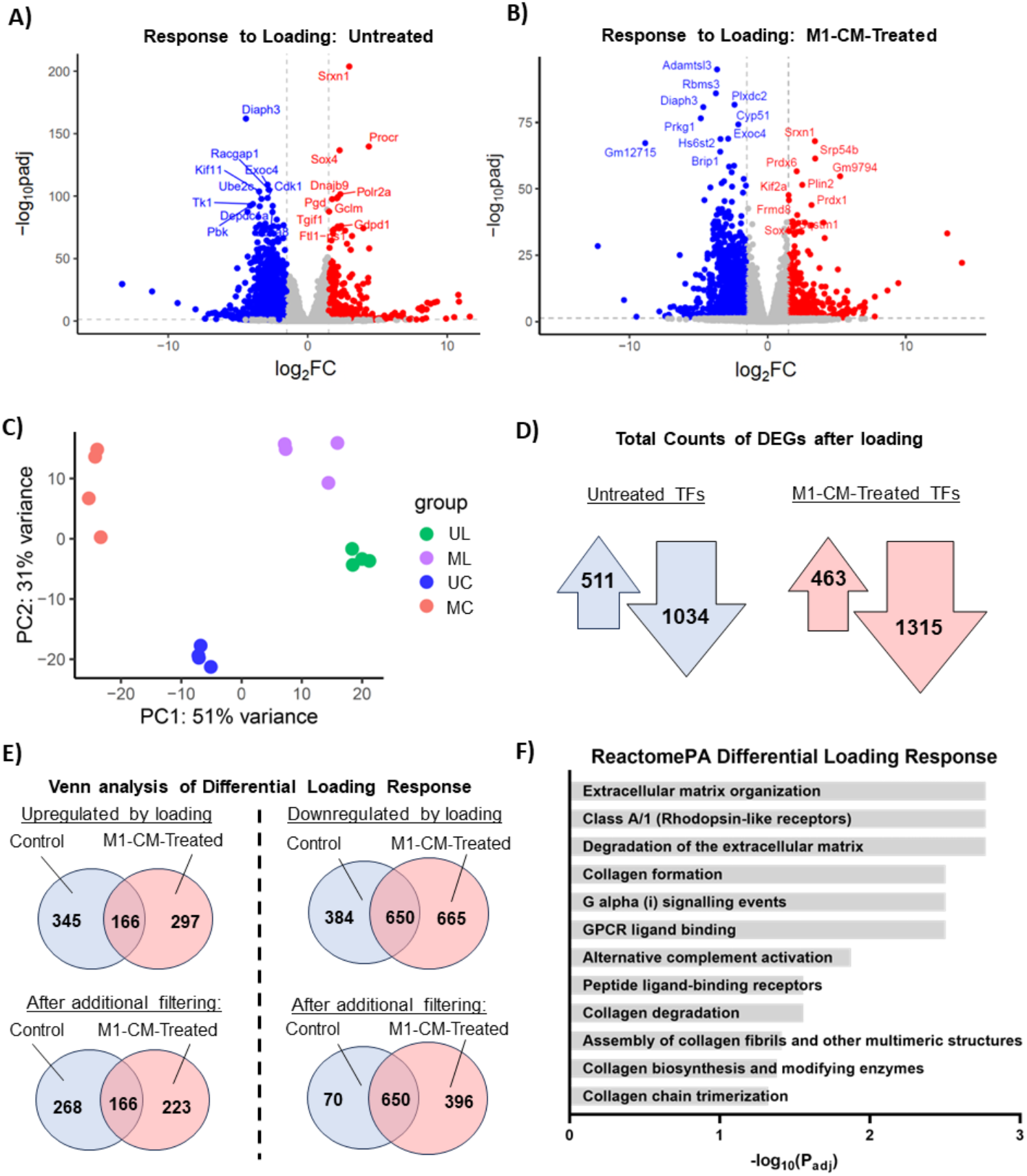
A) Volcano plot of DEGs in untreated TFs in response to loading. B) Volcano plot of DEGs in M1-CM-treated TFs in response to loading. C) PCA of all four TF groups: untreated control (UC), untreated loaded (UL), M1-CM-treated unloaded (MC), and M1-CM-treated loaded (ML). D) Total counts of significantly downregulated and upregulated genes in response to loading in both untreated (blue arrows) and M1-CM-treated (pink arrows) TFs. E) Overview of Venn analysis to identify genes regulated differently by loading in TFs after treatment with M1-CM. F) Significantly upregulated pathways among genes differentially regulated by loading depending on treatment condition as identified by Venn analysis.

### The NF-κβ and JAK/STAT pathways differentially mediated TF responses to M1-CM

The NF-κβ and JAK/STAT pathways were both upregulated in response to M1-CM, as shown by the pathway analysis of the RNA sequencing in Figure 2C. We therefore sought to determine whether these pathways were necessary for M1-CM-mediated TF responses. To inhibit NF-κβ, an inhibitor of the IKK-2 complex in the NF-κβ pathway was used a range of concentrations from 1uM to 10uM (Figure S1). To inhibit JAK/STAT, the same concentrations were tested using the JAK/STAT inhibitor Ruxolitinib. Figure 7 compares the effects of each inhibitor on the TF gene expression response to M1-CM, at the same dose of 2.5uM. Both inhibitors significantly suppressed the M1-CM-induced *Il6* and *Sod3* responses. However, only NF-κβ inhibition significantly suppressed M1-CM-induced *Ccl2*, and *Cxcl1*. TF responses Meanwhile, only JAK/STAT inhibition suppressed the M1-CM-induced *Mmp13* TF response.The effectiveness of NF-κβ inhibition increased with increasing concentration for all genes except *Mmp13*. By contrast, JAK/STAT inhibition did not appear to be more effective at higher doses in the range of concentrations tested (Figure S1). Effectiveness of suppression of M1-CM response by NF-κβ and JAK/STAT inhibitors was confirmed by Western Blot assays for phosphorylated p65 and phosphorylated STAT1, respectively (Figure S2).

**Figure 7:**
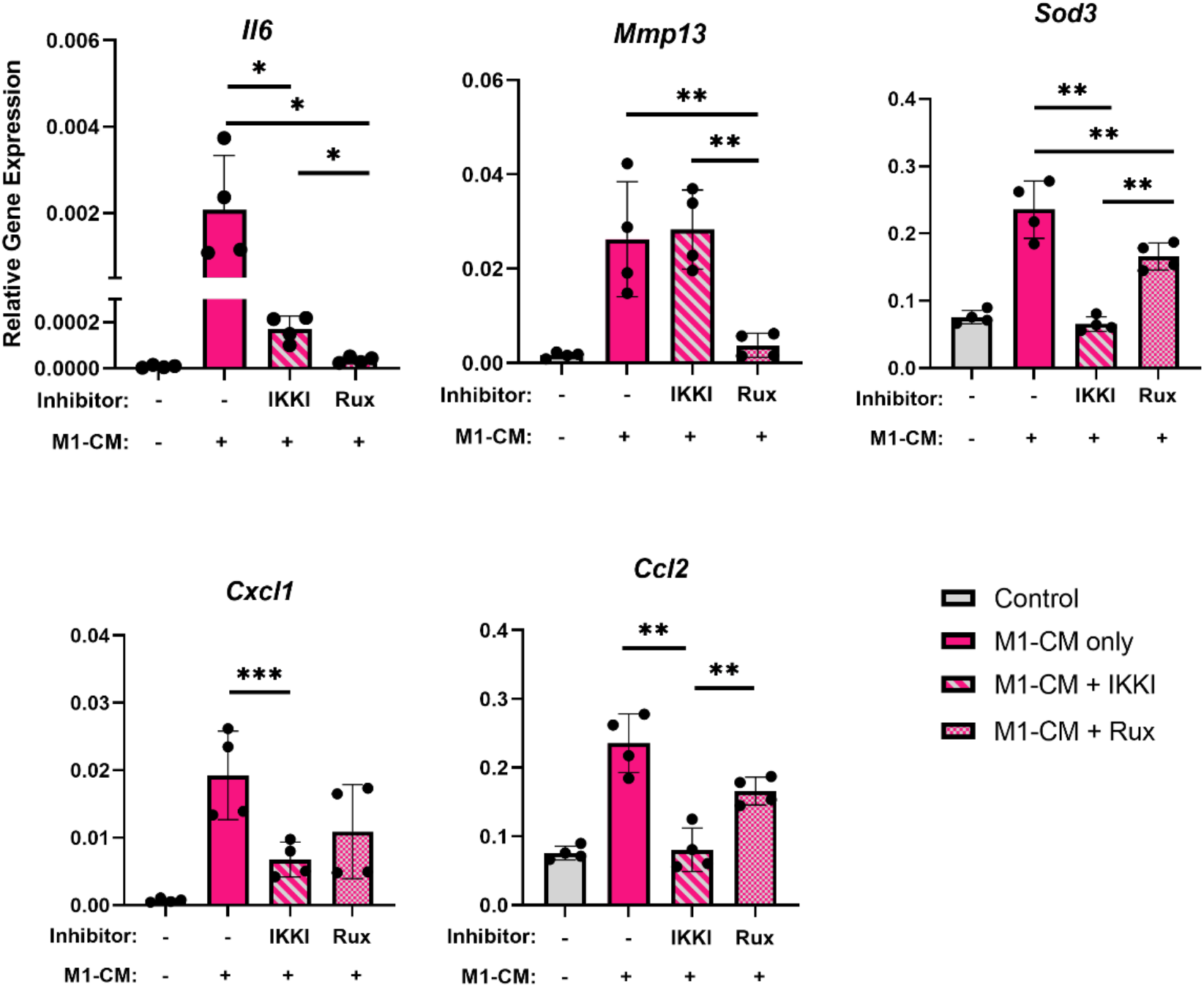
Comparison of effect of NF-κβ inhibitor (IKKI) vs. JAK/STAT inhibitor (Rux) on gene expression response of TFs to M1-CM-treatment. Data is shown as mean +/- SD, with individual points representing biologically independent samples. Statistical significance was calculated with a one-way ANOVA with Holm-Sidak adjustment for multiple comparisons between three groups (M1-CM only, M1-CM + IKKI, and M1-CM + Rux). *indicates p<0.05, **indicates p<0.01, ***indicates p<0.001.

Increasing concentrations of NF-κβ inhibitor led to corresponding decreases in cell confluence in the presence of M1-CM. To further investigate this qualitative observation, live/dead staining and the LDH assay were performed. Live/dead imaging confirmed lower numbers of live cells in the presence of NF-κβ inhibitor, with the lowest cell viability seen at the highest inhibitor concentration tested (Figure 8A). Interestingly, this effect was only observed in the presence of M1-CM, and was not seen under normal culture conditions. There was no observable effect of JAK/STAT inhibition on cell viability over the same range of concentrations. An LDH assay supported these results, showing significantly increased LDH compared to M1-CM-treated TFs in only the higher concentration NF-κβ inhibition groups (2.5uM and 10uM IKKI) (Figure 8B).

**Figure 8:**
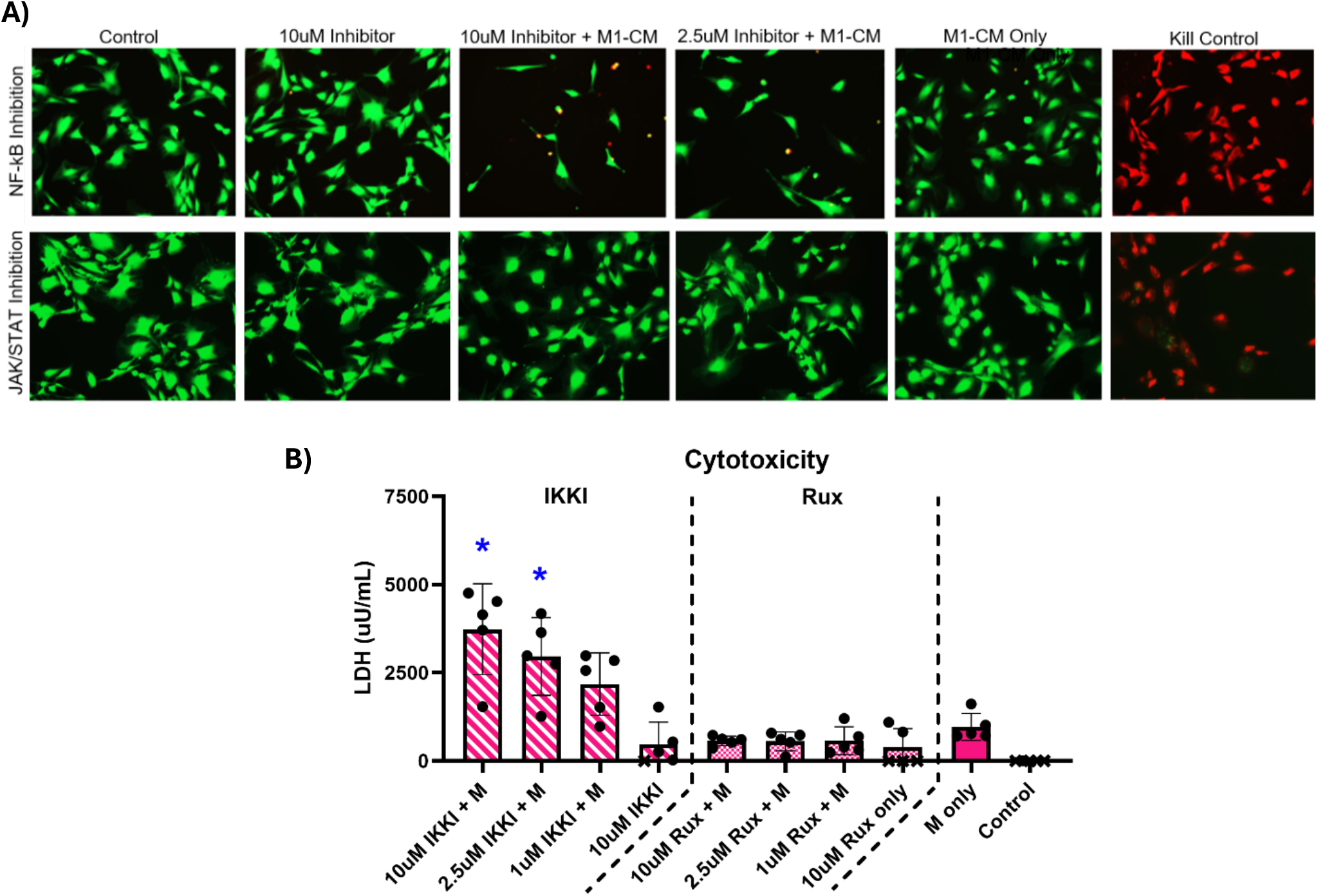
A) Live (green) / dead (red) stain to visualize differences in cell confluence. B) LDH assay for quantification of cytotoxicity of each inhibitor. Data is shown as mean +/- SD, with individual points (dot or x) representing biologically independent samples. M indicates M1-CM-treated, x indicates LDH levels below detection limit. IKKI indicates NF-κβ inhibitor, Rux indicates JAK/STAT inhibitor. Statistical significance was calculated with a one-way ANOVA with Holm-Sidak adjustment for multiple comparisons between M1-CM only and all M1-CM plus inhibitor groups.

### Mathematical modeling of time-dependent inflammatory suppression of tendon mechanotransduction

To better understand the interplay between inflammatory signaling and mechanical loading, we developed a simplified mathematical model that captures the suppression of mechanotransduction under inflammatory conditions (Figure 9). The model demonstrated that inflammatory factors decay exponentially following M1-CM treatment, with a half-life of approximately 7 hours (Figure 9A). Under control conditions, mechanical loading induced a robust ∼5-fold increase in IL-6 expression that closely matched our experimental data (Figure 9B, blue lines and dots). However, in the presence of M1-CM, IL-6 expression was constitutively elevated (∼8-fold) and showed minimal additional response to mechanical loading, with model predictions accurately capturing the experimental observations (Figure 9B, red lines and dots). Sensitivity analysis revealed that the inflammatory suppression parameter *K*_*inf*_ strongly influenced the final IL-6 levels, with lower values resulting in more complete suppression of mechanical responses (Figure 9C). The temporal analysis of mechanical responsiveness, calculated as the ratio of loaded to unloaded conditions, showed that control TFs maintained a consistent ∼5-fold mechanical response throughout the 24-hour period, while M1-CM-treated TFs exhibited near-complete loss of mechanosensitivity (Figure 9D), providing a quantitative framework for understanding how inflammation disrupts normal tendon mechanotransduction.

**Figure 9:**
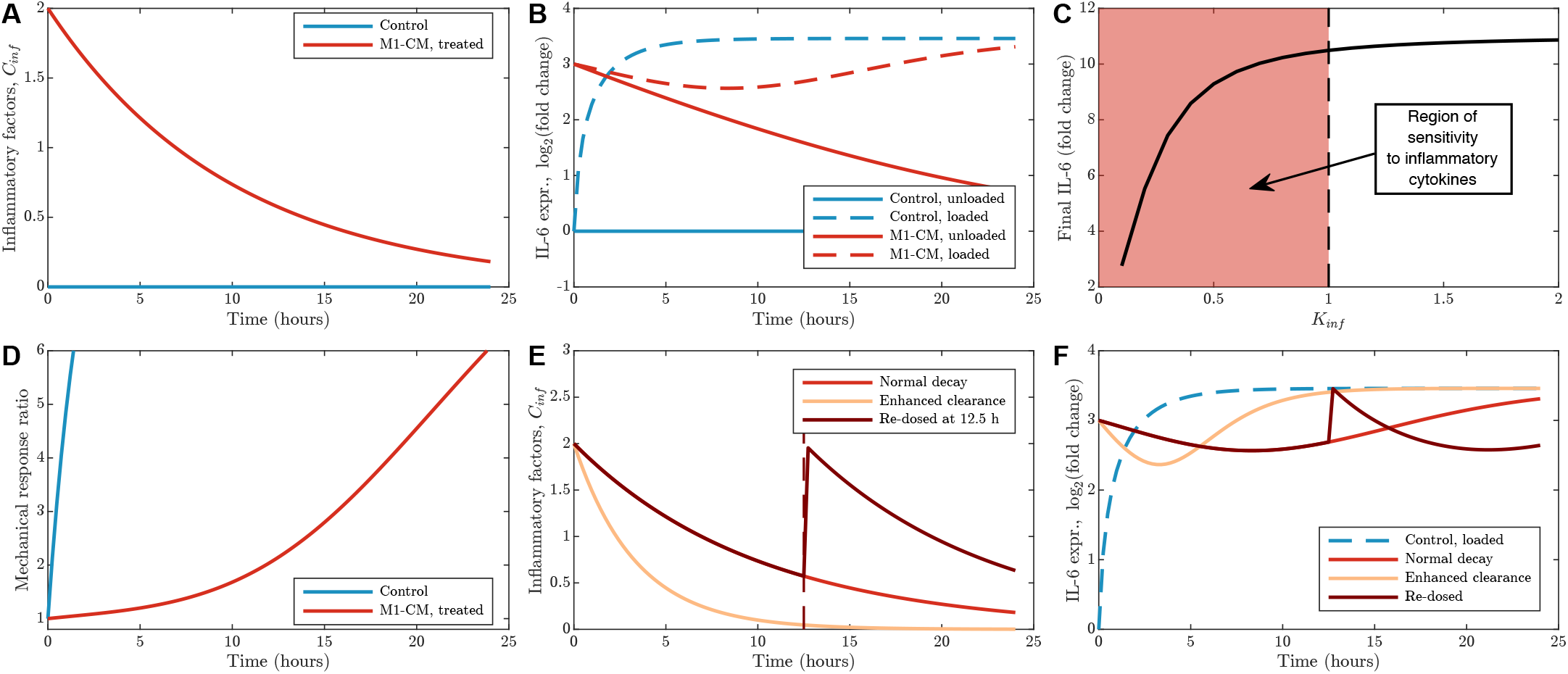
Mathematical model of IL-6 mechanotransduction under inflammatory conditions and therapeutic modulation strategies. (A) Temporal dynamics of inflammatory factors (*C*_*inf*_) showing decay from initial M1-CM stimulation (red line) compared to control conditions (blue line) over 24 hours. (B) Model predictions for IL-6 expression shown as log<sub>2</sub> fold change relative to unloaded control. Under control conditions (blue), mechanical loading induces a ∼10-fold increase in IL-6 (solid vs. dashed lines). Under inflammatory conditions (red), IL-6 expression is constitutively elevated (∼8-fold) with minimal additional response to mechanical loading, demonstrating inflammatory suppression of mechanotransduction. (C) Sensitivity analysis showing how the inflammatory suppression parameter (*K*_*inf*_) affects final IL-6 levels after 24 hours of loading under inflammatory conditions. Lower *K*_*inf*_ values result in stronger suppression of mechanical responses. M1-CM is a strong mediator of IL-6 levels for *K*_*inf*_ in the shaded region. (D) Time course of mechanical responsiveness, calculated as the ratio of loaded to unloaded IL-6 expression. Control conditions (blue) maintain robust mechanical responsiveness (∼5-fold), while inflammatory conditions (red) show near-complete loss of mechanical responsiveness throughout the 24-hour period. (E) Therapeutic modulation scenarios showing three different inflammatory dynamics: normal decay (red, *k*_*deg*_ = 0.1 h^−1^), enhanced clearance representing anti-inflammatory therapy (green, *k*_*deg*_ = 0.3 h^−1^), and re-dosing at 12.5 hours simulating repeated inflammatory insult (magenta, dashed line indicates re-application of M1-CM). (F) IL-6 responses under therapeutic scenarios with mechanical loading. Enhanced clearance (green) partially restores mechanical responsiveness by 24 hours, approaching control loaded levels (blue dashed). Re-dosing (magenta) maintains elevated IL-6 and prevents recovery of mechanosensitivity. Model parameters: *k*_*deg*_ = 0.1 h^−1^, *k*_*basal*_ = 0.1, *k*_*mech*_ = 2.0, *k*_*str*_= 0.5 h^−1^, *K*_*σ*_= 0.2, *K*_*inf*_ = 0.5, *n* = *m* = 2.

## Discussion

This study demonstrated that paracrine signals from M1 macrophages induce a proinflammatory response in TFs, NF-κβ and JAK/STAT signaling modulates that response, and TF inflammatory responses are altered by mechanical loading. Characterization of M1-CM and its effect on TFs revealed a robust inflammatory response in TFs, including the upregulation of over 500 genes and increased secretion of several cytokines. Additionally, quantification of cytokine secretion revealed that macrophages were able to induce an inflammatory profile in TFs that was distinct from the M1-CM inflammatory profile, implying a tendon tissue-specific response. Both JAK/STAT and NF-κβ were necessary for the response to M1-CM, and each pathway was responsible for different downstream responses to inflammation in TFs. TF responses to loading were altered by the presence of inflammatory stimuli, with more genes being downregulated by loading in M1-CM than under control conditions. These findings lay the foundation for further studies investigating inflammatory modulation and mechanical loading parameters for improved tendon healing.

M1 macrophages were chosen as the source of inflammation in our model system due to their dominant role in the tendon post-injury response ^11,36^. M1 macrophages can induce ECM degradation, and secrete a variety of cytokines such as IL-1β and TNFα that activate the immune response of surrounding cells ^46^. M1 macrophages can also directly influence fibroblast behavior by inducing upregulation of their pro-inflammatory gene expression and regulating their proliferation ^47,48^. In our model, the extent to which this single cell type was able to activate such a wide variety of pathways implied a process more representative of the inflammatory environment *in vivo* than prior experiments, which typically used a single cytokine such as IL-1β to induce inflammation ^49,50^. In addition, the response was achieved with levels of cytokines in the M1-CM that were orders of magnitude smaller than what is typically used *in vitro* to induce inflammation ^51–55^. Other studies have shown that the effects of a combination of two or three cytokines are much more potent than the same total quantity of a single cytokine ^24^. By using macrophage media, we were effectively treating TFs with a wider range of cytokines than other studies typically use, and at a more physiologically relevant dose. It has been shown previously that in the synovial fluid of the knee, for example, concentrations of cytokines after injury are on the order of pg/mL rather than ng/mL, which is much closer to what we measured in M1-CM ^56^.

Another intriguing finding of this study was that TFs produced cytokines at different proportions and magnitudes than M1 macrophages. Production of IL-6, for example, was 10 times higher in media from TFs stimulated with M1-CM than in M1-CM alone, in contrast to IFNγ production, which was not produced by TFs at meaningful levels. Additionally, the levels of chemokines were much higher in TFs + M1-CM supernatant than in M1-CM alone, suggesting that the macrophages themselves may not be the primary cell type responsible for continuing to recruit additional immune cells to the site post-injury ^57–59^. In fact, immune cell recruitment may be controlled in large part by resident fibroblasts rather than macrophages. This may be a mechanism by which resident TFs coordinate tissue repair, with M1 macrophages inducing the initial response, but TFs guiding the response thereafter.

Pathway analysis of RNAseq data showed that M1-CM significantly upregulated a variety of pathways in TFs, including TNFα, TLR, IL-17, NF-κβ, and JAK/STAT. These pathways have all been implicated in tendon injury, but the roles of NF-κβ and JAK/STAT in tendon healing have recently been of particular interest ^23,24^. Thus, these were investigated further in inhibition studies. In contrast, fewer pathways were significantly downregulated by M1-CM; one pathway of note was mTOR, which was previously identified as critical in TF mechanosensing ^56^. This result is consistent with our overall finding that the presence of M1-CM altered TF mechanoresponses. Other pathways that were downregulated in response to M1-CM were related to metabolism and stress-response. Detrimental effects of inflammation on metabolic function have been well documented in the context of several diseases, including rheumatoid arthritis, cardiovascular diseases, and neurological disorders, where inflammation has been shown to be linked to mitochondrial dysfunction ^60–62^. Specifically, treatment with IL1β and TNFα in human chondrocyte cells reduced activity of complex I in mitochondria and suppressed production of ATP ^63^. In tendon, altered metabolism has been observed in injured tendon, however the mechanisms and consequences of this are not fully understood and require further investigation to determine the role of metabolism in tendon pathology ^64^.

To determine the extent to which the NF-κβ and JAK/STAT pathways were necessary for the TF response to M1-CM, inhibition of these pathways in the presence of M1-CM was performed. These pathways were chosen due to our initial findings with M1-CM effects on TFs and due to recent literature that has highlighted their importance in tendon injury and their promise in therapeutic interventions to improve healing ^24,65^. While both pathways contributed to the TF responses to inflammation, they were each responsible for distinct downstream responses. *Mmp13* upregulation by M1-CM, for example, was only suppressed by JAK/STAT inhibition, and was unchanged with NF-κβ inhibition. This suggests that ECM degradation is driven more by JAK/STAT signaling than by NF-κβ signaling ^66^. Additionally, *Ccl2* and *Cxcl1* were unaffected by JAK/STAT inhibition, but were suppressed in the presence of the NF-κβ inhibitor. Therefore, NF-κβ likely plays a larger role in TF-mediated recruitment of immune cells than JAK/STAT ^57,58^. Future studies could investigate macrophage migration in response to TFs with and without NF-κβ inhibitor to confirm this hypothesis. Finally, NF-κβ inhibition suppressed the expression of *Sod3*, which encodes an enzyme that regulates oxidative stress by catalyzing the dismutase of superoxide, and which was significantly upregulated in response to M1-CM treatment ^44^. Therefore, the protective mechanism of TFs against stress was suppressed by NF-κβ inhibition. The opposite effect was observed with JAK/STAT inhibition, which suggests that JAK/STAT can suppress some of the detrimental effects of inflammation without suppressing all protective responses as well.

The relationship between NF-κβ inhibition and cell viability was an important finding in this study. While a role of NF-κβ in inhibiting apoptosis has been demonstrated in other processes, especially cancer treatment, this phenomenon has not been described in tendon ^67,68^. Additionally, because NF-κβ inhibition only reduced cell viability in the presence of inflammation, it is likely that NF-κβ is specifically playing a role in protecting against inflammation-mediated cell death. It is known that multiple pathways involved in inflammation can induce apoptosis in cells, including the TNF signaling pathway, which was shown to be upregulated in our RNA sequencing results ^69^. Additionally, TNFa was secreted in significant amounts by M1 macrophages. Thus, TNF signaling may induce apoptosis in TFs if NF-κβ is inhibited, which could be an important consideration for using NF-κβ targeted therapeutics in tendon injury. Future studies involving NF-κβ inhibition to improve tendon healing outcomes should investigate upregulation of apoptosis and monitor cellularity throughout healing.

Analysis of the genes that responded differently to loading under inflammatory conditions suggested changes in pathways involving extracellular matrix organization and G protein signaling. Interestingly, *Mmp13* and *Il6* were upregulated by loading under control conditions, but not under inflammatory conditions. In fact, *Il6* was significantly suppressed by loading after treatment with M1-CM. There are several potential explanations for this result. First, there were multiple genes upregulated by inflammation in the TFs which function as protective mechanisms against cell death and damage, including antioxidants like *Sod3*, and apoptosis-inhibiting pathways such as NF-κβ ^44,67,68^. Because these protective genes were upregulated by inflammation, they may prime TFs to suppress loading-induced stress responses such as increased *Il6* and *Mmp13*, in an effort to minimize damage. It is important to note, however, that the reduction in *Il6*, while significant, was not sufficient to bring *Il6* expression back down to the levels measured in untreated controls. Thus, loading may suppress the inflammatory response somewhat, but not completely.

RNA sequencing analysis revealed an altered loading response in hundreds of genes. When investigating these genes on a deeper level through Reactome Pathway Analysis, multiple pathways emerged related to ECM organization and collagen formation, which was supportive of our qPCR results for *Mmp13* and *Il6*. The “Extracellular matrix organization”, “Degradation of the extracellular matrix”, and “Collagen degradation” pathways were driven by several of the same genes, including *Mmp3* and *Mmp13*, which have been implicated in tendon disease ^70,71^. Most of the genes associated with these pathways were significantly upregulated in response to loading in untreated TFs, but not in M1-CM-treated TFs, which indicates that these ECM-related responses to loading are suppressed in the inflammatory environment. The “Collagen biosynthesis and modifying enzymes”, “Collagen formation”, and “Collagen chain trimerizations” pathways were driven by several overlapping genes as well, including *Col12a1, Col15a1, Col3a1, Col29a1*, and *Col8a2*. All of these genes were significantly downregulated in response to loading in M1-CM-treated TFs, but not in untreated TFs. This indicates that in the presence of inflammation, loading suppressed collagen synthesis. Considered together with the higher ECM degradation response to loading observed under control conditions, it is evident that inflammation overall suppressed the collagen turnover response to loading.

Additionally, “G alpha (i) signaling events”, “GPCR ligand binding”, and “Class A/1 (Rhodopsin-like receptors)” were identified in the Reactome Pathway Analysis. These pathways were driven by several shared genes, including *Hcar1, Npy*, and genes for several chemokines, including *Ccl20, Cxcl10*, and *Ccl11. Npy* was upregulated by loading in untreated TFs, but not in M1-CM-treated TFs, while *Hcar1* and the chemokines were downregulated by loading in the M1-CM-treated TFs, but not the control TFs. Both *Npy* and *Hcar1* have been shown to regulate calcium signaling and the ERK1/2 signaling cascade, known to be important in tendon mechanosensing ^72–74^. Additionally, chemokines have been shown to regulate intracellular calcium levels via GPCR signaling ^75^. Thus, the altered G protein signaling may a mechanism by which mechanotransduction is altered in TFs, but further studies directly measuring GPCR-related ERK/1 activation and calcium signaling would be necessary to confirm this hypothesis.

Our mathematical model, while simplified, reveals an intriguing possibility for therapeutic intervention that leverages interplay between inflammation and mechanical loading during tendon rehabilitation. The model predicts that enhanced inflammatory clearance (*k*_*deg*_ = 0.3 h−^1^) could lead to worse outcomes for chronic inflammation, with IL-6 levels approaching those of mechanically loaded controls by 24 hours (Figure 9E-F, green lines). This suggests that anti-inflammatory interventions might be most beneficial when timed to allow mechanical loading to stimulate appropriate healing responses. Conversely, the model demonstrates that re-dosing with inflammatory stimuli at 12.5 hours reduced elevated IL-6 levels transiently by reducing mechanosensitivity (Figure 9E-F, magenta lines). Although counterintuitive, this raises a testable hypothesis: carefully timed, periodic inflammatory stimulation might possibly enhance rehabilitation outcomes, allowing windows of mechanical responsiveness when inflammation subsides. This aligns with emerging evidence that some inflammation is necessary for proper healing, and suggests potential therapeutic strategies that might benefit from controlled inflammatory modulation rather than complete suppression. This motivates further modeling incorporating additional pathways and *in vivo* validation with periodic inflammatory dosing synchronized with rehabilitation protocols.

This work was not without limitations. First, the *in vitro* model system did not include a native ECM. Although changes in ECM synthesis, degradation, and reorganization can be postulated based on gene expression changes, ECM would need to be present to assess any net changes in ECM content at the protein level. Furthermore, attachments to the ECM play an important role in mechanotransduction. This has been shown to be the case for myofibroblasts in healing tendon, which are stimulated by inflammation to contract the ECM as part of the repair process. Second, the source of the inflammatory input was limited to macrophages. While this is the most abundant and persistent immune cell type in the early period after tendon injury, other cell types such as neutrophils and dendritic cells likely contribute as well. However, because macrophages are known to induce inflammation on their own in the context of tendon injury, and introducing other cell types would substantially increase the complexity and variability of the model system, it was assumed that the stimulating cytokines secreted by macrophages alone would be sufficient to largely recapitulate the inflammatory environment of the healing tendon. Third, this body of work focused largely on one-way inflammatory communication, specifically from macrophages to TFs. *In vivo*, any factors secreted by the tendon fibroblasts could affect the macrophages as well, and this this two-way communication was not fully modeled in this in vitro system. However, preliminary studies did not show any significant effect on macrophage phenotype resulting from the presence of the fibroblasts in co-culture, at least for the 24-hour time frame assessed. Additionally, the choice to isolate responses to one-way paracrine signaling had the benefit of illuminating a clearer interpretation of the TF-specific response to inflammation, which was the primary goal of this study.

In summary, TF responses to inflammation likely drive the progression of tendon pathology, as TFs produce large amounts of cytokines and significantly alter their gene expression in response to M1-CM. Inhibition of both the NF-κβ and JAK/STAT show promise in modulating the inflammatory environment of the injured tendon to improve healing, but further in vivo studies are necessary to determine how to translate this finding for improved healing outcomes. Furthermore, mechanical loading studies demonstrated that the application of cyclic strain in the presence of inflammation may serve to reduce ECM degradation processes and calm the inflammatory response in tendon, without suppressing it entirely.

## Acknowledgements

The study was funded in part by the NIH/NIAMS (R01 AR077793, R01 AR062947). RNA experiments were performed by the Columbia University Genome Center.

## References

1. Clayton, R. A. E. & Court-Brown, C. M. The epidemiology of musculoskeletal tendinous and ligamentous injuries. Injury 39, 1338–1344 (2008).

2. Ahmad, J., Repka, M. & Raikin, S. M. Treatment of myotendinous Achilles ruptures. Foot Ankle Int 34, 1074–1078 (2013).

3. Houshian, S., Tscherning, T. & Riegels-Nielsen, P. The epidemiology of Achilles tendon rupture in a Danish county. Injury 29, 651–654 (1998).

4. May, T. & Garmel, G. M. Rotator Cuff Injury. in StatPearls (StatPearls Publishing, Treasure Island (FL), 2024).

5. Frankewycz, B. et al. Achilles tendon elastic properties remain decreased in long term after rupture. Knee Surg Sports Traumatol Arthrosc 26, 2080–2087 (2018).

6. Thomopoulos, S. et al. The localized expression of extracellular matrix components in healing tendon insertion sites: an in situ hybridization study. J Orthop Res 20, 454–463 (2002).

7. Galatz, L. M., Ball, C. M., Teefey, S. A., Middleton, W. D. & Yamaguchi, K. The outcome and repair integrity of completely arthroscopically repaired large and massive rotator cuff tears. J Bone Joint Surg Am 86, 219–224 (2004).

8. Oh, L. S., Wolf, B. R., Hall, M. P., Levy, B. A. & Marx, R. G. Indications for rotator cuff repair: a systematic review. Clin Orthop Relat Res 455, 52–63 (2007).

9. Reito, A., Logren, H.-L., Ahonen, K., Nurmi, H. & Paloneva, J. Risk Factors for Failed Nonoperative Treatment and Rerupture in Acute Achilles Tendon Rupture. Foot Ankle Int. 39, 694–703 (2018).

10. Jiang, N., Wang, B., Chen, A., Dong, F. & Yu, B. Operative versus nonoperative treatment for acute Achilles tendon rupture: a meta-analysis based on current evidence. Int Orthop 36, 765–773 (2012).

11. Marsolais, D., Côté, C. H. & Frenette, J. Neutrophils and macrophages accumulate sequentially following Achilles tendon injury. J Orthop Res 19, 1203–1209 (2001).

12. Chartier, C. et al. Tendon: Principles of Healing and Repair. Semin Plast Surg 35, 211–215 (2021).

13. Altmann, N. et al. Interleukin-6 upregulates extracellular matrix gene expression and transforming growth factor β1 activity of tendon progenitor cells. BMC Musculoskelet Disord 24, 907 (2023).

14. Andersen, M. B., Pingel, J., Kjær, M. & Langberg, H. Interleukin-6: a growth factor stimulating collagen synthesis in human tendon. Journal of Applied Physiology 110, 1549–1554 (2011).

15. Ackermann, P. W., Domeij-Arverud, E., Leclerc, P., Amoudrouz, P. & Nader, G. A. Anti-inflammatory cytokine profile in early human tendon repair. Knee surg. sports traumatol. arthrosc. 21, 1801–1806 (2013).

16. Stauber, T. et al. Il-6 signaling exacerbates hallmarks of chronic tendon disease by stimulating reparative fibroblasts. eLife 12, RP87092 (2025).

17. Katsma, M. S. et al. The influence of chronic IL-6 exposure, in vivo, on rat Achilles tendon extracellular matrix. Cytokine 93, 10–14 (2017).

18. Chen, S. et al. Interleukin-6 Promotes Proliferation but Inhibits Tenogenic Differentiation via the Janus Kinase/Signal Transducers and Activators of Transcription 3 (JAK/STAT3) Pathway in Tendon-Derived Stem Cells. Med Sci Monit 24, 1567–1573 (2018).

19. Schulze-Tanzil, G. et al. The role of pro-inflammatory and immunoregulatory cytokines in tendon healing and rupture: new insights. Scandinavian Med Sci Sports 21, 337–351 (2011).

20. Chisari, E., Rehak, L., Khan, W. S. & Maffulli, N. The role of the immune system in tendon healing: a systematic review. British Medical Bulletin 133, 49–64 (2020).

21. Chisari, E., Rehak, L., Khan, W. S. & Maffulli, N. Tendon healing in presence of chronic low-level inflammation: a systematic review. British Medical Bulletin 132, 97–116 (2019).

22. Arvind, V. & Huang, A. H. Reparative and Maladaptive Inflammation in Tendon Healing. Front. Bioeng. Biotechnol. 9, 719047 (2021).

23. Golman, M. et al. Enhanced Tendon-to-Bone Healing via IKKβ Inhibition in a Rat Rotator cuff Model. The American journal of sports medicine 49, 780 (2021).

24. Avey, A. M., Devos, F., Roberts, A. G., Essawy, E. S. E. & Baar, K. Inhibiting JAK1, not NF-κB, reverses the effect of pro-inflammatory cytokines on engineered human ligament function. Matrix Biology 125, 100– 112 (2024).

25. Galatz, L. M. et al. Complete removal of load is detrimental to rotator cuff healing. J Shoulder Elbow Surg 18, 669–675 (2009).

26. Viidik, A. The Effect of Training on the Tensile Strength of Isolated Rabbit Tendons. Scandinavian Journal of Plastic and Reconstructive Surgery 1, 141–147 (1967).

27. Langberg, H., Rosendal, L. & Kjaer, M. Training-induced changes in peritendinous type I collagen turnover determined by microdialysis in humans. J Physiol 534, 297–302 (2001).

28. Ozone, K., Minegishi, Y., Oka, Y., Sato, M. & Kanemura, N. The Effects of Downhill Running and Maturation on Histological and Morphological Properties of Tendon and Enthesis in Mice. Biology (Basel) 12, 456 (2023).

29. Koskinen, S. O. A., Heinemeier, K. M., Olesen, J. L., Langberg, H. & Kjaer, M. Physical exercise can influence local levels of matrix metalloproteinases and their inhibitors in tendon-related connective tissue. J Appl Physiol (1985) 96, 861–864 (2004).

30. Reihmane, D., Jurka, A., Tretjakovs, P. & Dela, F. Increase in IL-6, TNF-α, and MMP-9, but not sICAM-1, concentrations depends on exercise duration. Eur J Appl Physiol 113, 851–858 (2013).

31. Yang, G.Im, H.-J. & Wang, J. H.-C. Repetitive mechanical stretching modulates IL-1β induced COX-2, MMP-1 expression, and PGE2 production in human patellar tendon fibroblasts. Gene 363, 166–172 (2005).

32. Archambault, J., Tsuzaki, M., Herzog, W. & Banes, A. J. Stretch and interleukin-1beta induce matrix metalloproteinases in rabbit tendon cells in vitro. J Orthop Res 20, 36–39 (2002).

33. McClinton, S. M., Heiderscheit, B. C., McPoil, T. G. & Flynn, T. W. Effectiveness of physical therapy treatment in addition to usual podiatry management of plantar heel pain: a randomized clinical trial. BMC Musculoskelet Disord 20, 630 (2019).

34. Desjardins-Charbonneau, A. et al. The Efficacy of Manual Therapy for Rotator cuff Tendinopathy: A Systematic Review and Meta-analysis. Journal of Orthopaedic & Sports Physical Therapy 45, 330–350 (2015).

35. Vicenzino, B., Branjerdporn, M., Teys, P. & Jordan, K. Initial changes in posterior talar glide and dorsiflexion of the ankle after mobilization with movement in individuals with recurrent ankle sprain. J Orthop Sports Phys Ther 36, 464–471 (2006).

36. Sugg, K. B., Lubardic, J., Gumucio, J. P. & Mendias, C. L. Changes in macrophage phenotype and induction of epithelial-to-mesenchymal transition genes following acute Achilles tenotomy and repair. J Orthop Res 32, 944–951 (2014).

37. Wang, J. H.-C., Thampatty, B. P.Lin, J.-S. & Im, H.-J. Mechanoregulation of gene expression in fibroblasts. Gene 391, 1–15 (2007).

38. Hong, Y. et al. Cell–matrix feedback controls stretch-induced cellular memory and fibroblast activation. Proc. Natl. Acad. Sci. U.S.A. 122, e2322762122 (2025).

39. Whitham, M. et al. Contraction-induced Interleukin-6 Gene Transcription in Skeletal Muscle Is Regulated by c-Jun Terminal Kinase/Activator Protein-1. Journal of Biological Chemistry 287, 10771– 10779 (2012).

40. Cahill, C. M. & Rogers, J. T. Interleukin (IL) 1β Induction of IL-6 Is Mediated by a Novel Phosphatidylinositol 3-Kinase-dependent AKT/IκB Kinase α Pathway Targeting Activator Protein-1. Journal of Biological Chemistry 283, 25900–25912 (2008).

41. Jäger, S., Handschin, C., St.Pierre, J. & Spiegelman, B. M. AMP-activated protein kinase (AMPK) action in skeletal muscle via direct phosphorylation of PGC-1α. Proc. Natl. Acad. Sci. U.S.A. 104, 12017– 12022 (2007).

42. Paschoud, S. et al. Destabilization of Interleukin-6 mRNA Requires a Putative RNA Stem-Loop Structure, an AU-Rich Element, and the RNA-Binding Protein AUF1. Molecular and Cellular Biology 26, 8228–8241 (2006).

43. Mishra, V. et al. IL-1β turnover by the UBE2L3 ubiquitin conjugating enzyme and HECT E3 ligases limits inflammation. Nat Commun 14, 4385 (2023).

44. Wert, K. J. et al. Extracellular superoxide dismutase (SOD3) regulates oxidative stress at the vitreoretinal interface. Free Radic Biol Med 124, 408–419 (2018).

45. Carow, B. & Rottenberg, M. E. SOCS3, a Major Regulator of Infection and Inflammation. Front Immunol 5, 58 (2014).

46. Mosser, D. M. & Edwards, J. P. Exploring the full spectrum of macrophage activation. Nat Rev Immunol 8, 958–969 (2008).

47. de la Durantaye, M., Piette, A. B., van Rooijen, N. & Frenette, J. Macrophage depletion reduces cell proliferation and extracellular matrix accumulation but increases the ultimate tensile strength of injured Achilles tendons. J Orthop Res 32, 279–285 (2014).

48. Manning, C. N. et al. Adipose-derived mesenchymal stromal cells modulate tendon fibroblast responses to macrophage-induced inflammation in vitro. Stem Cell Res Ther 6, 74 (2015).

49. Zhang, K., Asai, S., Yu, B. & Enomoto-Iwamoto, M. IL-1β Irreversibly Inhibits Tenogenic Differentiation and Alters Metabolism In Injured Tendon-Derived Progenitor Cells In Vitro. Biochem Biophys Res Commun 463, 667–672 (2015).

50. McClellan, A. et al. A novel mechanism for the protection of embryonic stem cell derived tenocytes from inflammatory cytokine interleukin 1 beta. Sci Rep 9, 2755 (2019).

51. Busch, F. et al. Resveratrol Modulates Interleukin-1β-induced Phosphatidylinositol 3-Kinase and Nuclear Factor κB Signaling Pathways in Human Tenocytes. J Biol Chem 287, 38050–38063 (2012).

52. Paterson, Y. Z., Rash, N., Garvican, E. R., Paillot, R. & Guest, D. J. Equine mesenchymal stromal cells and embryo-derived stem cells are immune privileged in vitro. Stem Cell Res Ther 5, 90 (2014).

53. Buhrmann, C. et al. Curcumin modulates nuclear factor kappaB (NF-kappaB)-mediated inflammation in human tenocytes in vitro: role of the phosphatidylinositol 3-kinase/Akt pathway. J Biol Chem 286, 28556–28566 (2011).

54. Riera, K. M. et al. Interleukin-1, tumor necrosis factor-alpha, and transforming growth factor-beta 1 and integrative meniscal repair: influences on meniscal cell proliferation and migration. Arthritis Res Ther 13, R187 (2011).

55. Mia, M. M., Boersema, M. & Bank, R. A. Interleukin-1β attenuates myofibroblast formation and extracellular matrix production in dermal and lung fibroblasts exposed to transforming growth factor-β1. PLoS One 9, e91559 (2014).

56. Mousavizadeh, R. et al. β1 integrin, ILK and mTOR regulate collagen synthesis in mechanically loaded tendon cells. Sci Rep 10, 12644 (2020).

57. Hsieh, S.-L. et al. MCP-1 controls IL-17-promoted monocyte migration and M1 polarization in osteoarthritis. Int Immunopharmacol 132, 112016 (2024).

58. Yadav, S. K. et al. Chemokine-triggered microtubule polymerization promotes neutrophil chemotaxis and invasion but not transendothelial migration. J Leukoc Biol 105, 755–766 (2019).

59. Anton, K., Banerjee, D. & Glod, J. Macrophage-Associated Mesenchymal Stem Cells Assume an Activated, Migratory, Pro-Inflammatory Phenotype with Increased IL-6 and CXCL10 Secretion. PLOS ONE 7, e35036 (2012).

60. Liu, J. T. et al. Mitochondrial function is altered in articular chondrocytes of an endemic osteoarthritis, Kashin-Beck disease. Osteoarthritis Cartilage 18, 1218–1226 (2010).

61. Dai, D.-F., Rabinovitch, P. S. & Ungvari, Z. Mitochondria and Cardiovascular Aging. Circ Res 110, 10.1161/CIRCRESAHA.111.246140 (2012).

62. Voloboueva, L. A. & Giffard, R. G. Inflammation, Mitochondria and the Inhibition of Adult Neurogenesis. J Neurosci Res 89, 1989–1996 (2011).

63. López-Armada, M. J. et al. Mitochondrial activity is modulated by TNFalpha and IL-1beta in normal human chondrocyte cells. Osteoarthritis Cartilage 14, 1011–1022 (2006).

64. Ackerman, J. E., Best, K. T., Muscat, S. N. & Loiselle, A. E. Metabolic Regulation of Tendon Inflammation and Healing Following Injury. Curr Rheumatol Rep 23, 15 (2021).

65. Lane, R. A. et al. The effects of NF-κB suppression on the early healing response following intrasynovial tendon repair in a canine model. J Orthop Res 41, 2295–2304 (2023).

66. Del Buono, A., Oliva, F., Osti, L. & Maffulli, N. Metalloproteases and tendinopathy. Muscles Ligaments Tendons J 3, 51–57 (2013).

67. Verzella, D. et al. Life, death, and autophagy in cancer: NF-κB turns up everywhere. Cell Death Dis 11, 1–14 (2020).

68. Bernal-Mizrachi, L., Lovly, C. M. & Ratner, L. The role of NF-{kappa}B-1 and NF-{kappa}B-2-mediated resistance to apoptosis in lymphomas. Proc Natl Acad Sci U S A 103, 9220–9225 (2006).

69. van Loo, G. & Bertrand, M. J. M. Death by TNF: a road to inflammation. Nat Rev Immunol 23, 289–303 (2023).

70. Corps, A. N., Curry, V. A., Buttle, D. J., Hazleman, B. L. & Riley, G. P. Inhibition of interleukin-1beta-stimulated collagenase and stromelysin expression in human tendon fibroblasts by epigallocatechin gallate ester. Matrix Biol 23, 163–169 (2004).

71. Sun, H. B. et al. Coordinate Regulation of IL-1β and MMP-13 in Rat Tendons Following Subrupture Fatigue Damage. Clin Orthop Relat Res 466, 1555–1561 (2008).

72. Lecat, S., Belemnaba, L.Galzi, J.-L. & Bucher, B. Neuropeptide Y receptor mediates activation of ERK1/2 via transactivation of the IGF receptor. Cell Signal 27, 1297–1304 (2015).

73. Popov, C. et al. Mechanical stimulation of human tendon stem/progenitor cells results in upregulation of matrix proteins, integrins and MMPs, and activation of p38 and ERK1/2 kinases. BMC Mol Biol 16, 6 (2015).

74. Nordström, F. et al. The lactate receptor GPR81 is predominantly expressed in type II human skeletal muscle fibers: potential for lactate autocrine signaling. Am J Physiol Cell Physiol 324, C477–C487 (2023).

75. Dixit, N. & Simon, S. I. Chemokines, selectins and intracellular calcium flux: temporal and spatial cues for leukocyte arrest. Front Immunol 3, 188 (2012).

